# Transient primary cilia mediate robust Hedgehog pathway-dependent cell cycle control

**DOI:** 10.1101/741769

**Authors:** Emily K. Ho, Anaïs E. Tsai, Tim Stearns

## Abstract

The regulation of proliferation is one of the primary functions of Hedgehog (Hh) signaling in development. Transduction of Hh signaling requires the primary cilium, a microtubule-based organelle that is necessary for several steps in the pathway (Corbit et al., 2005; Huangfu and Anderson, 2005; Huangfu et al., 2003; Liu et al., 2005; Rohatgi et al., 2007). Many cells only build a primary cilium upon cell cycle arrest in G0. In those proliferating cells that do make a cilium, it is a transient organelle, being assembled in G1 and disassembled sometime after, although exactly when is not well-characterized (Ford et al., 2018; Pugacheva et al., 2007; Wang and Dynlacht, 2018). Thus the requirement for primary cilia presents a conundrum: how are proliferative signals conveyed through an organelle that is present for only part of the cell cycle? Here we investigate this question in a mouse medulloblastoma cell line, SMB55, that requires cilium-mediated Hh pathway activity for proliferation (Zhao et al., 2015). We show that SMB55 cells are often ciliated beyond G1 into S phase, and the presence of the cilium determines the periods of Hh pathway activity. Using live imaging over multiple cell cycles, we define two windows of opportunity for Hh pathway activity, either of which is sufficient to effect cell cycle entry. The first is in the ciliated phase of the previous cell cycle, and the second is in G1 of the cell cycle in which the decision is made. We propose that the ability of cells to integrate Hh pathway activity from more than one cell cycle imparts robustness on Hh pathway control of proliferation and may have implications for other Hh-mediated events in development.

## Results

In the developing cerebellum, granule cell precursors require Hh signaling transduced by primary cilia for their proliferation, as do some cancerous medulloblastomas originating from this cell type (Barakat et al., 2013; Goodrich et al., 1997; Han et al., 2009; Spassky et al., 2008; Wechsler-Reya and Scott, 1999; Wong et al., 2009). We chose the mouse medulloblastoma line SMB55 for our experiments because the cells maintain this dependence on Hh pathway activity and cilia for growth in culture, which we confirmed (**Figure S1**) (Zhao et al., 2015, 2017). The Hh receptor Patched-1 (Ptch1) normally inhibits the activator Smoothened (Smo). SMB55 cells derive from a *Ptch1*^+/-^ mouse and have lost heterozygosity at the *Ptch1* locus (**Figure S2A**), so these cells have constitutively activated Smo and pathway activation independent of Sonic hedgehog (Shh) ligand. The Hh pathway in SMB55 cells can be inhibited by the Smo antagonist SANT-1, as shown by reduced expression of the transcription factor *Gli1* and cell cycle effector target genes *Ccnd1* and *Mycn* (**Figure 1A**) (Chen et al., 2002; Hatton et al., 2006; Kenney and Rowitch, 2000; Kenney et al., 2003; Oliver et al., 2003). We found that *Gli1* mRNA decreased after just 4 hours of treatment with 100 nM SANT-1 (**Figure 1B**). Thus, SANT-1 treatment provides a relatively rapidly-acting means to modulate Hh pathway activity.

**Figure 1:**
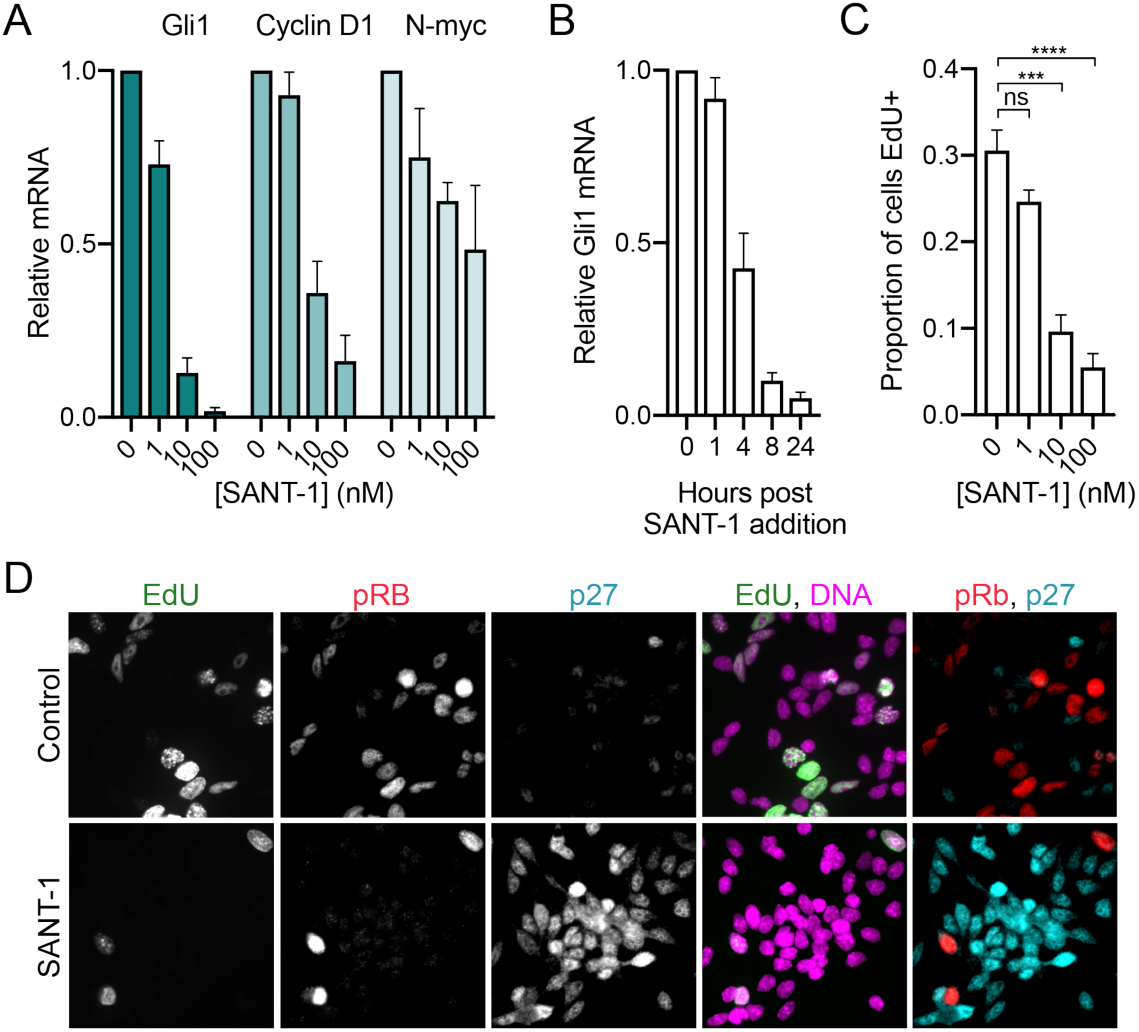
Hh pathway activity is required for SMB55 cell proliferation. (A) Hh pathway target mRNA after 24-hour inhibition with increasing concentrations of SANT-1. *Gli1, Ccnd1* and *Mycn* mRNAs were measured by RT-qPCR, normalized to GAPDH, and expressed relative to 0 nM controls. Mean+SEM, n=3 (B) Time course of *Gli1* mRNA after treatment with 100 nM SANT-1 for indiciated times. *Gli1* mRNA was measured by RT-qPCR, normalized to GAPDH, and expressed relative to controls at each time point. Mean+SEM, n=3 (C) Proportion of EdU^+^ SMB55 cells after 48-hour treatment with increasing concentrations of SANT-1 (EdU added for last two hours). Mean+SEM from 3 independent experiments, n ≥250 cells in each. Significance determined by one-way ANOVA and Tukey’s post hoc test. (D) SMB55 cells labeled with EdU and stained with antibodies against pRb and p27 after 48-hour treatment with 100 nM SANT-1. Scale bar: 10 μm See also Figure S1, S2.

We next assayed the nature of the reduction in SMB55 cell proliferation after Hh pathway inhibition. After 48-hour SANT-1 treatment, we labeled cells with the nucleotide analog EdU for 2 hours to measure the proportion of SMB55 cells in S phase. SANT-1 treatment significantly reduced the proportion of EdU^+^ cells in a dose-dependent manner (**Figure 1C, D**), whereas it had no effect on cells whose growth does not require the Hh pathway (**Figure S2B**). We then stained for p27 and phospho-Rb (pRb), markers of cell cycle arrest and commitment respectively.

Most SANT-1-treated cells were p27^+^ and pRb^-^, consistent with G0 arrest (**Figure 1D**). SANT-1-treated cells also maintained expression of Atoh1, a marker of cerebellar precursors, indicating arrest rather than differentiation (**Figure S2C**). Therefore, SMB55 cells require Hh pathway activity for cell cycle entry at G1/S.

We next assessed SMB55 cell ciliation during the cell cycle. Staining for primary cilia in an asynchronous population using the cilia membrane marker Arl13b revealed that most cells had a cilium (**Figure 2A**). We then expressed lentivirally-produced eYFP-PCNA in these cells to mark S-phase cells with nuclear PCNA puncta. Many S-phase cells were ciliated (**Figure 2A**), contrary to the widely-held view that the primary cilium is limited to G0/G1 phase, but consistent with the recent results of Ford, et al. (2018). To investigate further, we arrested cells for 24 hours in S phase using 2 mM hydroxyurea (HU), a ribonucleotide reductase inhibitor, or in G2 phase using 10 µM RO-3306, a Cdk1 inhibitor. Cilia quantification revealed no difference in the proportion of ciliated S phase-arrested cells compared to the asynchronous control population, whereas the proportion of ciliated G2-arrested cells was significantly reduced (**Figure 2B**). Therefore, SMB55 cells typically have a cilium during the cell cycle, including S phase, but lose the cilium by G2 phase.

**Figure 2:**
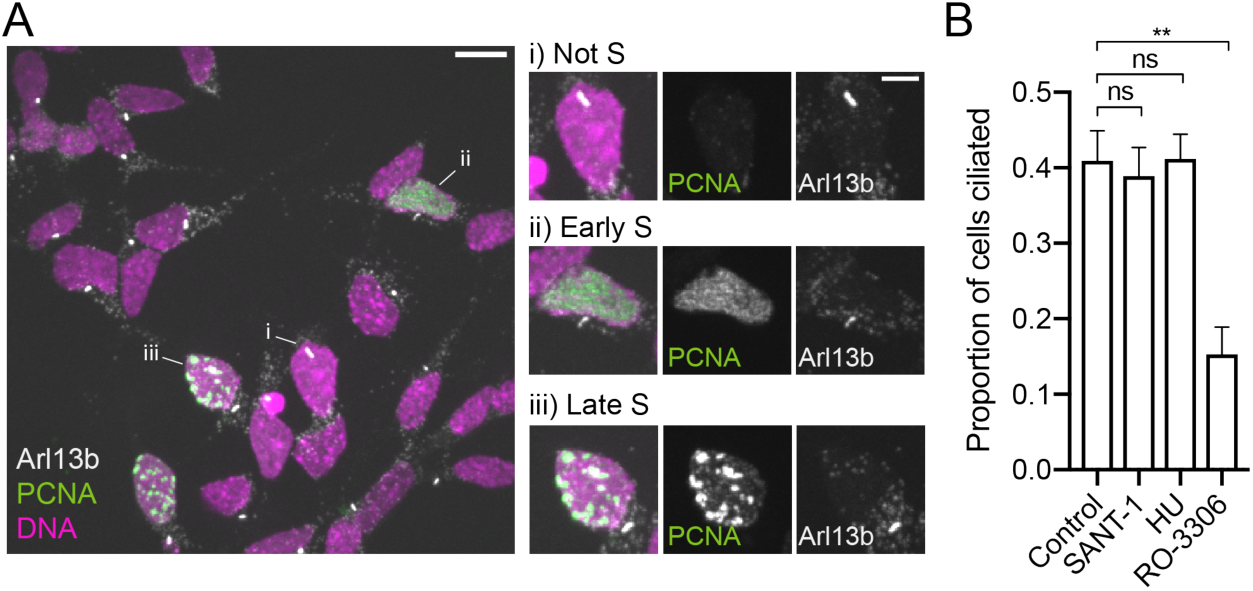
Presence of cilia on SMB55 cells during the cell cycle. (A) Asynchronous SMB55 population expressing eYFP-PCNA and stained for Arl13b and DNA. Insets highlight ciliated SMB55 cells (i) not in S, (ii) in early S, and (iii) in late S, distinguished by nuclear PCNA foci. Scale bar: 10 μm. Inset: 5 μm (B) Proportion of cells ciliated after 24-hour treatment with 100 nM SANT-1, 2 mM HU, or 10 μM RO-3306 compared to an asynchronous population. Mean+SEM from 3 independent experiments, n ≥100 cells in each. Significance determined by one-way ANOVA with Tukey’s post hoc test.

Because the mechanisms of ciliary signaling have largely been studied in G0, we tested whether S-phase cilia are competent for Hh signal transduction. Here we used NIH3T3 cells, a highly Hh-responsive mouse fibroblast line, to allow temporal control over activation of the pathway. We treated NIH3T3 cells with activating concentrations of Shh or SAG (a Smo agonist) for just 2 hours, reasoning that most cells would remain in the same cell cycle phase for the duration of the treatment. We identified ciliated S-phase cells (PCNA^+^, Arl13b^+^) and measured ciliary Smo to assess pathway activation. For all experimental conditions, there was no significant difference in ciliary Smo in S-phase compared to non-S-phase cilia; both populations had increased ciliary Smo upon pathway activation (**Figure 3A, 3A’**). We found similar results with SMB55 cells, comparing the constitutively activated state to SANT-1 inhibition. Ciliary Smo intensity was similar in S-phase compared to non-S-phase SMB55 cells; both populations had high Smo in control cells and reduced Smo in SANT-1-treated cells (**Figure 3B, 3B’**). We then asked whether presence of the cilium is temporally correlated with pathway activation within a cell cycle. We measured *Gli1* mRNA by RT-qPCR in SMB55 cells arrested in S phase with HU for 24 hours and found that these cells had high *Gli1* mRNA, comparable to control cells (**Figure 3C**). The half-life of *Gli1* mRNA is significantly shorter than 24 hours (Noubissi et al., 2009). Therefore the Hh pathway must be transcriptionally active in ciliated S-phase SMB55 cells. In contrast, RO-3306-arrested SMB55 cells, which typically do not have a cilium (see **Figure 2B**), had low *Gli1* mRNA, comparable to SANT-1-treated cells (**Figure 3C**); this effect is likely specific to the cell cycle stage, as RO-3306 had no effect on Hh signaling itself (**Figure S3**). Thus, S-phase cilia are competent for Hh signal transduction, and in SMB55 cells, which have constitutively active Smo, it is presence or absence of a cilium that determines Hh pathway activity during the cell cycle.

**Figure 3:**
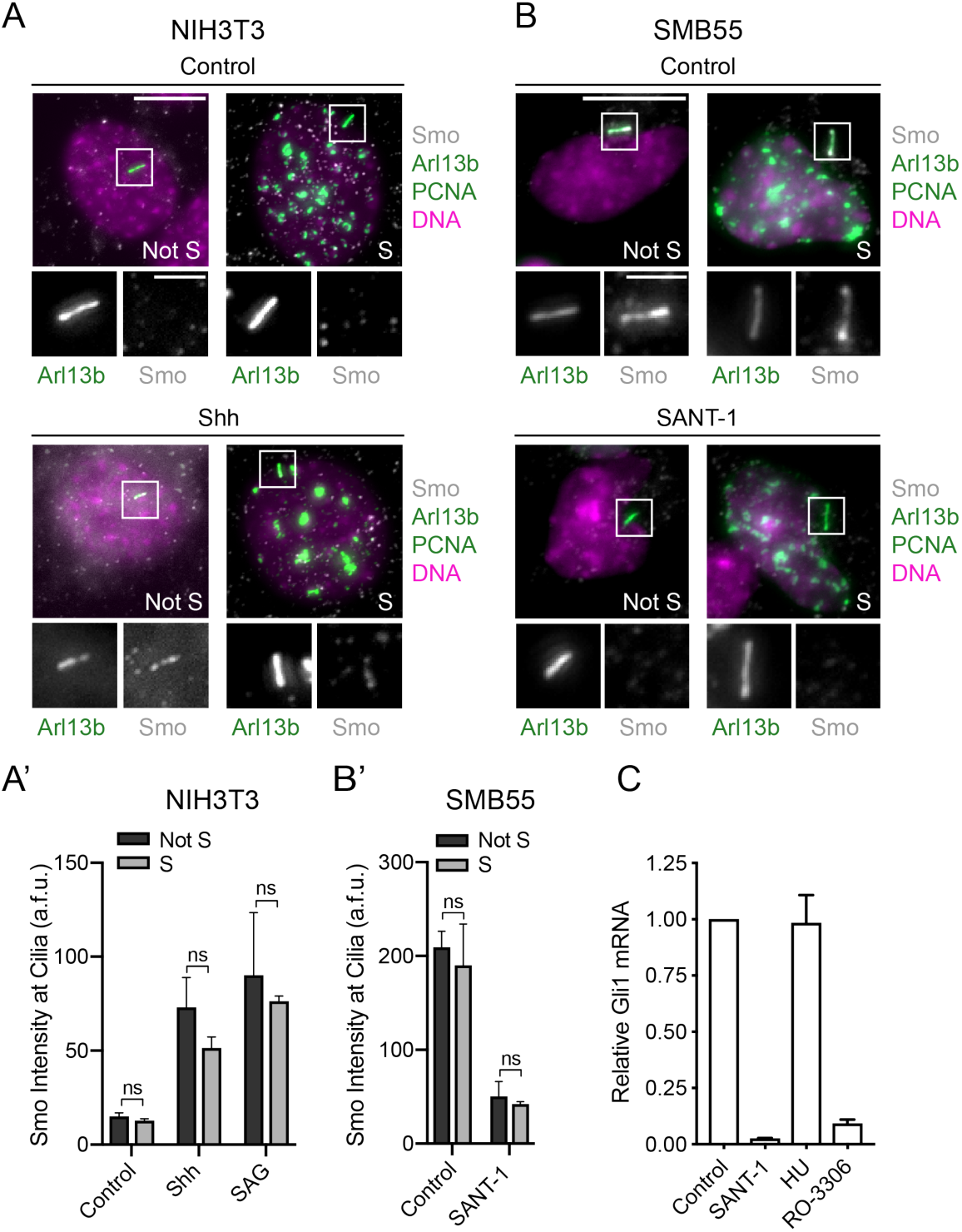
Cilium presence determines Hh pathway activity during the cell cycle. (A-B) Smo intensity, an indicator of pathway activity, in S-phase and non-S-phase cilia of (A) eYFP-PCNA NIH3T3 cells after 2-hours pathway stimulation with Shh or SAG and (B) eYFP-PCNA SMB55 cells untreated or after 24-hour pathway inhibition with SANT-1. Cells were stained for Smo and cilia (Arl13b), and PCNA was used to identify S-phase cells. Scale bar: 10 μm, Inset: 3 μm. (A’, B’) Quantification of Smo intensity (Mean+SEM) at cilia from 3 independent experiments, n=30 cells in each. No significant difference in any condition between S-phase and non-S-phase cilia by two-way ANOVA with Tukey’s post hoc test. (C) *Gli1* mRNA in SMB55 cells treated with 100 nM SANT-1, 2 mM HU, or 10 μM RO-3306 for 24 hours. *Gli1* mRNA amount was measured by RT-qPCR, normalized to GAPDH and expressed relative to the asynchronous control. n=3 See also Figure S3.

Having defined the periods of Hh pathway activity during the SMB55 cell cycle, we next asked when signaling is required to inform the G1/S cell cycle entry decision. Using live imaging, we tracked individual, asynchronously-growing cells through multiple cell cycles, using an approach analogous to that used to study the yeast cell cycle (Hartwell et al., 1974). Addition of SANT-1 to the asynchronous population at the beginning of imaging allowed us to observe the effect of Hh pathway inhibition on cells starting at all points in the cell cycle. Through long-term imaging, we could then determine the outcome of the cell cycle entry versus exit decision by assessing whether each cell entered the ensuing mitosis (M_2_) (**Figure 4A**). For clarity, we refer to the cell cycle in which we are assessing the entry decision as the “decision cell cycle” and the one previous to that as the “previous cell cycle.” Because visualization of individual cells in dense neurospheres, as SMB55 are typically grown, was not feasible, we performed 48-hour phase-contrast imaging in 2D cultures. We confirmed that SMB55 cells under these conditions still required Hh pathway activity by showing that most of the SANT-1-treated cells were not cycling at the end of the imaging period (**Figure 4B**).

**Figure 4:**
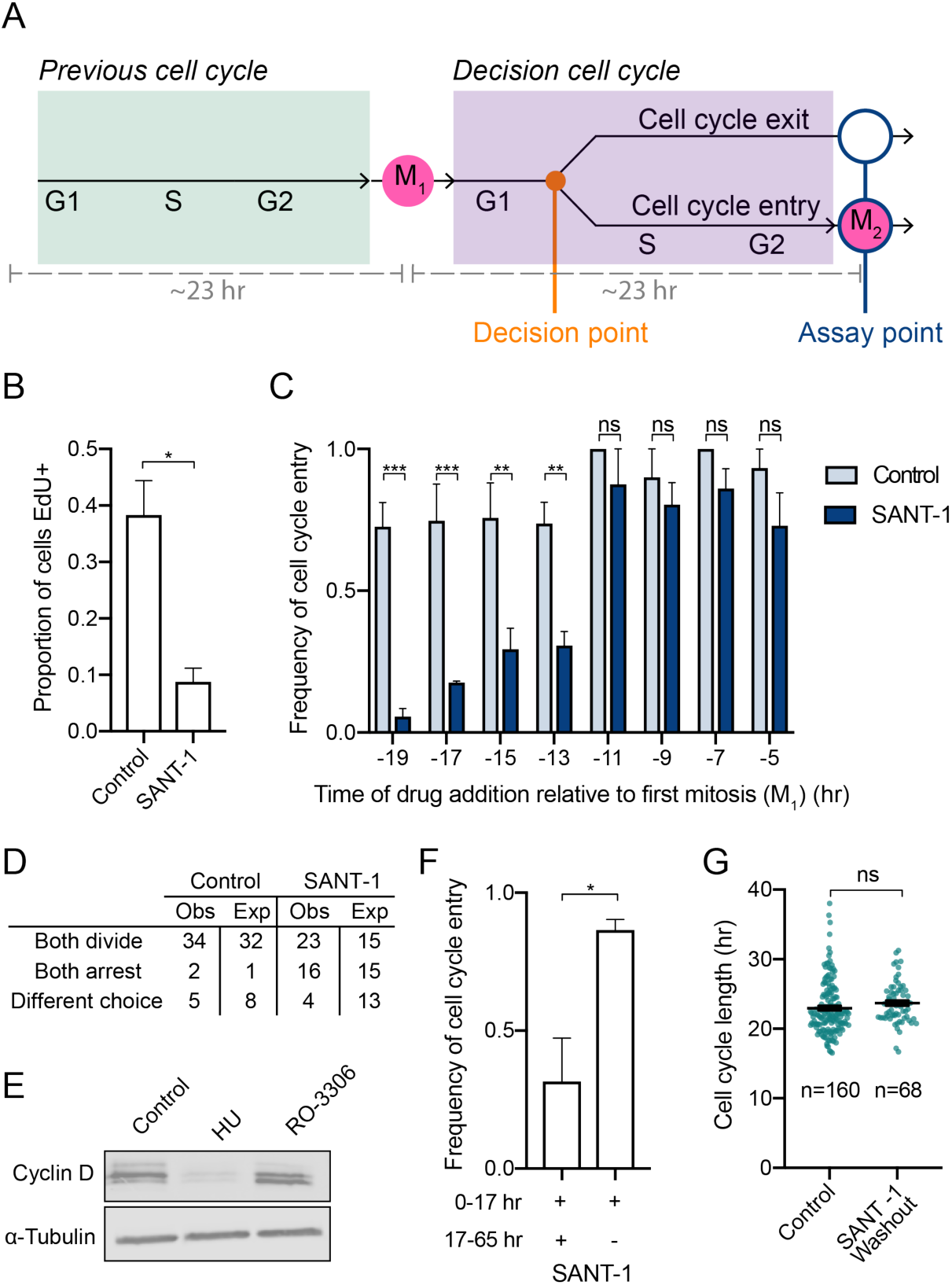
Hh pathway activity in the previous cell cycle or the decision cell cycle is sufficient for cell cycle entry. (A) Diagram of the cell cycle decision, highlighting the decision point and the mitotic assay point in reference to the “previous” and “decision” cell cycles. (B) Proportion of SMB55 cells incorporating EdU following 48-hour SANT-1 treatment in 2D imaging conditions. Mean+SEM from 3 independent experiments, n ≥200 cells in each. (C) Effect of SANT-1 treatment in the previous cell cycle on SMB55 cell cycle entry. Based on the time of drug addition relative to M_1_, the frequency of cells progressing to M_2_ is shown, representing the frequency of cell cycle entry. Mean+SEM from 3 independent experiments, n=3-20 cells in each. (D) Division decisions made by sister cells, comparing those observed to those expected if sister cells behaved independently. The expected distributions for control and SANT-1-treated cells were based on cell cycle entry frequencies in (C). The distribution of SANT-1-treated cell outcomes was significantly different from the expected distribution (*χ*^2^= 6.481, df =2, *p*=0.0391); the control cell distribution was not (*χ*^2^=1.086, df =2, *p*=0.58). (E) Cyclin D expression in SMB55 cells treated with 2 mM HU or 10 μM RO-3306 for 24 hours. Cyclin D was measured by western blot with α-tubulin as a loading control. (F) Effect of SANT-1 washout at M_1_ on SMB55 cell cycle entry. The frequency of cell cycle entry after 17-hour SANT-1 treatment and washout is shown. Mean+SEM from 3 independent experiments, n=10-40 cells in each. (G) Cell cycle length of SMB55 cells following SANT-1 washout. Each point represents the time between M_1_ and M_2_ of a single cell, for all control (C) and SANT-1-washout (F) cells. Each dot represents one cell, Mean±SEM also shown. Significance determined by (B, F-G) unpaired Student’s t test, (C) two-way ANOVA and Tukey’s post hoc test. See also Video S1.

To analyze the image sequences, we identified the first division (M_1_) after SANT-1 addition for each cell. Based on the time of M_1_ relative to that of SANT-1 addition, we determined how long the Hh pathway had been inhibited in the previous cell cycle. We then tracked each cell from M_1_ onward to determine whether it entered the decision cell cycle, and thus continued to mitosis M_2_ (**Video S1**). Remarkably, SMB55 cells treated with SANT-1 for up to 11 hours before M_1_ entered the cell cycle as frequently as control cells (**Figure 4C**). Only when the pathway was inhibited for more than 12 hours before M_1_ did the cells fail to enter the decision cell cycle. Given that 4-hour SANT-1 treatment is sufficient to inhibit the Hh pathway (see **Figure 1B**), the pathway would not have been active in the decision cell cycle in any of the tracked cells. Thus, Hh pathway activity in G1 of the decision cell cycle is not necessary for cell cycle entry; signaling in the previous cell cycle is sufficient.

If Hh pathway activity in the previous cell cycle determines cell cycle entry, we would predict that sister cells derived from the same parental cell would make the same decision more often than two independently-derived cells. We compared the observed distribution of sister-cell decisions to the expected distribution according to the cell cycle entry frequencies measured in Figure 4C. For SANT-1-treated cells, the decisions differed significantly from the expected distribution, suggesting that sister-cell decisions are not independent and supporting our model (**Figure 4D**). We also would predict that Cyclin D protein would begin to accumulate in the previous cell cycle when the Hh pathway is active. Accordingly, western blotting showed low Cyclin D in HU-arrested cells but high Cyclin D in RO-3306-arrested cells, consistent with degradation in S and accumulation in G2 (**Figure 4E**), as observed previously by Yang et al. (2006).

Since SMB55 cells typically bear a cilium both in G1-S of the previous cell cycle and G1 of the decision cell cycle, we investigated whether pathway activity in G1 of the decision cell cycle would also be sufficient for cell cycle entry. We treated SMB55 cells with SANT-1 for 17 hours and then washed out the inhibitor, assaying cell cycle entry in those cells at M_1_ at the time of washout. Significantly more cells entered the cell cycle after washout compared those remaining in SANT-1 past M_1_ (**Figure 4F**). These cells also had similar cell cycle lengths to control cells, arguing against a need to spend a longer time accumulating sufficient proliferative signal in G1 (**Figure 4G**). Therefore, Hh pathway activity in either the previous cell cycle or in G1 of the decision cell cycle is sufficient for cell cycle entry.

## Discussion

A conundrum of cell signaling in vertebrates is that some pathways require the cilium, but the cilium is present only transiently during the cell cycle of dividing cells. Our results demonstrate that SMB55 medulloblastoma cells, which require Hh pathway activity for proliferation, make a cilium during G1 and maintain that cilium through S phase. We confirmed that SMB55 cell proliferation requires the primary cilium and showed that Hh pathway activation is not limited to G1/G0, but is possible whenever cells are ciliated. Thus SMB55 cells have two opportunities for Hh pathway activity prior to cell cycle entry: G1-S of the previous cell cycle, and G1 of the cell cycle in which they are making the entry decision. Our results reveal that Hh pathway activity in either of these signaling windows is sufficient for cell cycle entry. The presence of two sufficient signaling windows supports a model in which Hh pathway activity is integrated over the cell cycle rather than measured at a single decision point, and the cell cycle entry decision is made when a threshold of pathway activity is reached. This result is reminiscent of the Hh pathway’s control of differentiation in the neural tube, where integration of signal over time determines differential target gene expression (Dessaud et al., 2007).

The presence of two sufficient signaling windows also reveals that the cell cycle entry decision is robust to perturbations in pathway activity beyond those we observed due to changes in ciliation state during the cell cycle. Interestingly, robustness is a well-established feature of Hh morphogen gradient interpretation, imparted by mechanisms such as negative feedback and temporal adaptation (Barkal and Leibler, 1997; Ribes and Briscoe, 2009). The need for robustness is usually considered to derive from fluctuations in Hh ligand, which certainly would be relevant for proliferation in non-cancerous contexts. However, neither cilium assembly nor disassembly is a synchronous process (Anderson and Stearns, 2009; Mirvis et al., 2019), suggesting that ciliation itself may also be a source of variability in the signaling level a given cell can attain. New tools to manipulate ciliation are necessary to directly test how these fluctuations affect proliferation.

Our results indicate that SMB55 cells can reach the threshold required for cell cycle entry as early as halfway through the previous cell cycle (approximately S phase). We define this point as the restriction point for Hh pathway activity (Pardee, 1974) and note that recent work has similarly placed the restriction point for non-ciliary mitogen signaling in the previous cell cycle (Spencer et al., 2013). A restriction point in the previous cell cycle requires that some element of the signaling pathway persist into the decision cell cycle, which in this case requires persistence through G2, when the cells we studied, and most cells, typically lack a cilium. This persistence in the absence of continued signaling could occur through stable downstream effects of the signaling, as has been described previously for Cyclin D mRNA (Yang et al., 2017). Consistent with this, we showed that expression of Cyclin D can occur outside of the ciliated phases of the cell cycle. However, additional mechanisms, such as epigenetic changes (Shi et al., 2014), may also be involved.

Here we have used SMB55 cells as a model of Hh-pathway dependent proliferation and shown that because of ciliation during the cell cycle and pathway activation during ciliated portions, these cells have multiple windows of opportunity for Hh pathway activation that together impart robustness on the cell-cycle entry decision. Our work highlights the importance of considering cilia dynamics during the cell cycle when investigating cilium-dependent signaling events. It will be particularly interesting to test the implications of these findings on Hh’s role as a morphogen during developmental decisions.

## Materials and Methods

### Lead Contact and Materials Availability

Further information and requests for resources and reagents should be directed to and will be fulfilled by the Lead Contact, Tim Stearns.

### Experimental Model and Subject Details

#### SMB55 cell culture

SMB55 cells were a gift of R. Segal (Dana Farber Cancer Institute, Harvard Medical School). SMB55 cells were cultured in DMEM/F-12 without L-Glutamine (Corning) supplemented with 2% B-27 without retinoic acid (Gibco) and 1% penicillin/streptomycin (HyClone). Cells were maintained as neurospheres in 3D culture in untreated 96 well plates (Corning), changing media every 2 days. Cells were dissociated every 6 days with Accutase (Sigma-Aldrich) and plated at 150,000 cells/mL. All experiments, except for the live imaging experiments, were performed in 3D culture. Cells were frequently tested to confirm that growth was dependent on Hh pathway activity using SANT-1 treatment and EdU incorporation as described below. SMB55 cells were genotyped to confirm the absence of the wild-type *Ptch1* allele using primers recognizing the wild-type (Ptch-WT3: 5’-CTGCGGCAAGTTTTTGGTTG- 3’, Ptch-WT4: 5’-AGGGCTTCTCGTTGGCTACAAG-3’) and mutant alleles (NeoF3: 5’-TGGGGTGGGATTAGATAAATGCC-3’, PtchR3: 5’- TGTCTGTGTGTGCTCCTGAATCAC-3’) as described in (Goodrich et al., 1997). Genotyping PCR was performed using Taq polymerase on genomic DNA extracted from SMB55 cells using ZymoBead™ Genomic DNA kit (Zymo). Genomic DNA extracted from *Ptch1*^*-/-*^ mouse embryonic fibroblasts and a wild-type mouse were used as a control for the mutant and wild-type alleles respectively. All cells were grown in a humidified incubator at 37°C, 5% CO_2_.

#### Other cell culture

NIH3T3 cells were obtained from ATCC (RRID:CVCL_0594) and cultured in DMEM (Corning) plus 10% FBS (Gemini). To maximize ciliation of S-phase cells, NIH3T3 cells were split from dense plates and plated in dense (but not confluent) conditions. To induce ciliation in NIH3T3 cells, confluent cultures were starved in 0.5% FBS for 24 hours. HEK293T/17 cells were obtained from ATCC (RRID:CVCL_1926) and cultured in DMEM plus 10% CCS (Hyclone). All cells were grown in a humidified incubator at 37°C, 5% CO_2_.

### Method Details

#### Antibodies used

Primary antibodies used to immunofluorescence: rabbit anti-phospho-Rb Ser807/811, clone D20B12 (1:1000, Cell Signaling, RRID:AB_11178658), mouse IgG1 anti-p27^Kip^, clone 57 (1:500, BD), mouse IgG1 anti-polyglutamylated tubulin, clone GT335 (1:500, Adipogen RRID:AB_2490210), mouse IgG2a anti-Arl13b, clone N295B/66 (1:1000, Neuromab RRID:AB_2341543), rabbit anti-Smo (1:500, described previously (Rohatgi et al,. 2007)), mouse IgG1 anti-gamma tubulin, clone GTU-88 (1:5000, Sigma-Aldrich RRID:AB_532292), mouse IgG1 anti-Atoh1(Math1) (supernatant 1:10, Developmental Studies Hybridoma Bank RRID:AB_10805299), rabbit anti-Cep164 (1:500, described previously in (Lee et al., 2014), rabbit anti-Cyclin D (1:1000, Millipore RRID:AB_10562237), mouse IgG1 anti-alpha tubulin, clone DM1A (1:10,000, Sigma-Aldrich RRID:AB_310035). For immunofluorescence, AlexaFluor conjugated secondary antibodies (Thermo-Fisher) were diluted 1:1000. For western blotting, IRDye conjugated secondary antibodies (LI-COR) were diluted 1:20,000.

#### EdU incorporation assays

SMB55 cells were plated at 150,000 cells/mL in an untreated 96 well plate. NIH3T3 cells were plated on coverslips at 40,000 cells/well of a 24 well plate. Drug was added at the time of plating. At 24 hours, fresh drug was added and SMB55 cells were dissociated with Accutase. At 48 hours, Click-iT EdU (10 µM) was added to cell culture media and incubated for 2 hours. After the labeling period, the EdU was washed out and SMB55 cells were allowed to adhere to laminin-coated coverslips for approximately 3 hours before fixation, still in the presence of drug.

#### Hedgehog pathway manipulation

The Smo antagonist SANT-1 (100 nM, Millipore) and agonist SAG (100 nM, Enzo Life Sciences) were used to inhibit or activate the Hh pathway for the indicated length of time. DMSO was used as a vehicle control in all assays involving SANT-1 or SAG. Human Sonic hedgehog (Shh) carrying two isoleucine residues at the N-terminus and a hexahistidine tag at the C-terminus was expressed in *Escherichia coli* Rosetta2(DE3)pLysS cells (Novagen) and purified by immobilized metal-affinity chromatography followed by gel-filtration chromatography as described previously (Bishop et al., 2009). The purified protein was used in signaling assays at 100 nM.

#### Cell cycle arrest

Cell cycle arrest in SMB55 cells was performed by adding Hydroxyurea (2mM, Sigma-Aldrich) or RO-3306 (10 µM, AdipoGen) at the time of plating, treating for 24 hours, and then collecting cells for downstream analysis. DMSO was used as a vehicle control in all assays involving RO-3306.

#### Immunofluoresence and Click-it labeling

For fluorescence microscopy experiments, cells were fixed on poly-D lysine-coated (Sigma) glass #1.5 coverslips (Electron Microscopy Sciences). For SMB55 cells, these coverslips were coated overnight with 10 µg/mL laminin (Gibco) dissolved in DMEM/F-12 without L-Glutamine. The SMB55 neurospheres were allowed to adhere to the coverslips for 3-24 hours before fixation. NIH3T3 cells were grown on coverslips for the duration of the experiment. Cells were either fixed in 4% formaldehyde in PBS for 15 minutes at room temperate or 100% methanol for 15 minutes at −20°C. Fixed cells were washed 3 times for 5 minutes in PBS and then blocked in 1% Normal Donkey Serum (Jackson Immuno Research), 0.1% Triton X-100 in PBS for 30 minutes. Primary antibody was diluted in blocking solution and incubated on coverslips for 1 hour at room temperature. Coverslips were washed with PBS, and then AlexaFluor conjugated secondary antibody (Invitrogen) was diluted 1:1000 in blocking solution and incubated on coverslips for 1 hour at room temperature. Coverslips were washed with PBS again and incubated with DAPI (Sigma-Aldrich) for 10 minutes. After a final wash, coverslips were mounted with Mowiol 4-88 (Calbiochem) in glycerol containing 1,4,-diazobicycli-[2.2.2] octane (DABCO, Sigma-Aldrich) antifade. The click reaction between EdU and Alexafluor-594 azide was performed with the Click-iT 594 Labeling Kit according to the manufacturer’s instructions (Invitrogen). Cells were imaged on either a wide-field Axiovert 200M microscope (Zeiss) with PlanApoChromat 63x/1.4 NA objective and OrcaER CCD camera (Hamamatsu) or a AxioObserver microscope (Zeiss) with a confocal spinning disk head (Yokogawa, Japan), PlanApoChromat 63x/1.4 NA objective, and a Cascade II:512 EMCCD camera (Photometrics). All images were acquired with Micro-Manager software (Edelstein et al., 2014). Images of primary cilia were taken in 4-5 µm stacks (Wide field: 1 µm slices, Confocal: 0.5 µm slices) to ensure all cilia in the field of view were included.

#### Lentivirus production and infection

To make recombinant lentivirus, HEK293T/17 cells were cotransfected with the respective transfer vector and second-generation lentiviral cassettes (packaging vector psPAX2 (Addgene RRID:Addgene_12260) and envelope vector pMD2.G (Addgene RRID:Addgene_12259)) using 1 mg/mL polyethylenimine (PEI; Polysciences). The medium was changed 24 hours after transfection, and viral supernatant was harvested after an additional 48 hours. To concentrate the lentivirus, supernatant was filtered through a 0.45 µm filter and viral particles were pelleted by centrifugation (2:20 hours, 20°C, 50,000 x g). The viral pellet was then resuspended in PBS at 4°C overnight. For lentiviral infection of SMB55 cells, concentrated lentivirus was added to cells at the time of plating in the presence of 4 µg/mL Hexadimethrine bromide (Polybrene, Sigma-Aldrich) for an infection efficiency of about 80%. Media was changed 24 hours post infection. For infection of NIH3T3 cells, concentrated lentivirus was added to cells 24 hours after plating in the presence of 8 µg/mL Polybrene for an infection efficiency of about 80%. Media was changed 24 hours post infection. For experiments involving lentiviral expression of pTRIP eYFP-PCNA (gift of J. Ferrell), infected cells were selected in 400 µg/mL G418 (Gemini) beginning 2 days post infection for 2 weeks. A clonal NIH3T3 eYFP-PCNA line was isolated by limiting dilution and used for all experiments.

#### shRNA knockdown

For shRNA knockdown, the oligos targeting Mouse Cep164 (5’-GGTGATCTTTACTATTTCA-3’) and Firefly Luciferase (5’-TGAAGTCTCTGATTAAGTA-3’) were cloned into the HpaI and XhoI restriction sites in pSICOR (Addgene RRID:Addgene_11579) (Ventura et al., 2004). shRNAs targeting the same region of Firefly Luciferase have been used previously (Premsrirut et al., 2011). The empty, pSICOR plasmid was used as an additional control. Knockdown efficacy and effect on ciliation were tested by immunofluorescence at 4 days post infection. Proliferation was measured by a 2-hour EdU incorporation 4 days post infection.

#### RT-qPCR

For measurement of mRNA transcript amount by RT-qPCR, 300,000 SMB55 cells were treated for the indicated amount of time and then collected by centrifugation. Cells were resuspended in 1 mL Trizol reagent (Invitrogen) and total RNA was extracted according to the manufacturer’s instructions. The reverse transcription reaction was performed using the Maxima First Strand Synthesis kit (Thermo Scientific) with 185 ng of RNA. The product of this reaction was then assessed by qPCR in the Applied Biosystems Fast 7500 using the Taqman™ Gene Expression Master Mix (Applied Biosystems) and multiplexed Taqman™ probes. All reactions were performed in triplicate. Target gene values were normalized to GAPDH levels from the same reaction and expressed relative to control samples using the ΔΔCt method. The following mouse probes (Applied Biosystems) were used: GAPDH VIC-MGB (Mm99999915), Gli1 FAM-MGB (Mm00494645_m1), Cyclin D1 FAM-MGB (Mm00432359_m1), N-Myc FAM-MGB (Mm00476449_m1).

#### Immunoblotting

For immunoblotting, 1×10^6^ SMB55 cells were treated for 24 hours and then collected by centrifugation. Cells were lysed by shaking in 50 µL RIPA buffer (25 mM Tris 7.4, 150 mM NaCl, 2% NP-40, 0.25% Sodium Deoxycholate, 1 mM EDTA, 1 mM DTT) supplemented with 1 mg/ml each leupeptin, pepstatin, and chymostatin, and 1 mM phenylmethylsulfonyl fluoride for 1 hour at 4°C. Soluble lysate was then isolated by centrifugation (1 hour, 4°C, 16,100 x g). Protein concentration was measured by BCA and then 30 µg of protein was boiled in sample buffer and run on an SDS-PAGE gel. After transfer to nitrocellulose membrane (BioRad), membranes were blocked in 5% milk in TBST for 30 min, and then incubated in primary antibody diluted in TBST overnight. Membranes were washed and then incubated in secondary antibody solution for 1 hour. IRDye conjugated secondary antibodies (LI-COR) were diluted 1:20,000 in TBST. Membranes were washed again and scanned on a LI-COR Odyssey scanner.

#### Live Imaging

For live imaging of SMB55 cells, 525,000 SMB55 cells were plated as neurospheres in 3D culture. After 2 days of growth, spheres were plated in fresh media on 35 mm Fluorodish glass imaging dishes (World Precision Instruments) that had been coated with poly-D lysine and then laminin as described for immunofluorescence microscopy. SANT-1 was added at the time of plating. Cultures were imaged by phase-contrast microscopy on a Keyence BZ-X700 all-in-one fluorescence microscope with a Plan Fluor 20x/0.45 NA Ph1 objective (Nikon) for 48 hours at 5-minute intervals in a humidified, temperature-controlled (37°C), and CO_2_-controlled (5%) chamber (Tokai Hit). For washout experiments, cells on imaging dishes were treated for 16.5 hours with SANT-1 beginning at the time of plating. To washout the SANT-1, three 5-minute washes were performed with fresh media. In the washout condition, this media contained DMSO only as a vehicle control. In the non-washout condition, this media still contained SANT-1. The imaging was started at the end of this procedure and the washout was considered to begin at 17 hours post SANT-1 addition. We selected a 17-hour SANT-1 treatment prior to M_1_ because without washout this condition largely prevented cell cycle entry (see Figure 4C). Cultures were then imaged for 48 hours. At the end of all live imaging experiments, cells were labeled with EdU for 2 hours before fixation to confirm the effect on SANT-1 on the population’s proliferation before performing analysis of single cells.

#### Live imaging analysis

All live imaging analysis was performed in FIJI. To analyze image sequences for the live imaging experiment, all mitotic events in the first 4-19 hours after drug addition were identified. The frame containing cytokinesis was used to define the time of mitosis relative to drug addition. Daughter cells of each mitotic event were then tracked manually through the image sequences. If the cell divided, the time of cytokinesis was recorded and the cell was considered to have entered the cell cycle. If the cell was tracked to the end of the image sequence without dividing, it was counted as not dividing and thus not entering the cell cycle. Cells that left the field of view, were not able to be tracked with certainty, or had an aborted or multipolar mitosis were not included in the analysis. For sister cell analysis, only pairs in which both cells could be defined as entering the cell cycle or not were included. In the washout experiment, cells born from mitotic events only in the first 2 hours after washout were tracked and analyzed, in order to capture cells in which SANT-1 was washed out as close to M_1_ as possible. Tracking was performed similarly as above, except that cells were considered to not enter the cell cycle if they did not divide within 30 hours. Cell cycle length was determined by the cytokinesis-cytokinesis time in cells that did enter the cell cycle and divide.

#### Image quantification

All image quantification was performed in FIJI. To quantify EdU incorporation, the number of EdU^+^ nuclei was expressed relative to the total number of nuclei (identified by DAPI). Proportion of cells ciliated, Smo intensity, and Cep164 intensity were quantified from maximum projections of a 5 µm z-stack (1 µm slices) taken on the Axiovert. The proportion of cells ciliated was determined by counting the number of cilia in a field of view and dividing that count by the number of nuclei. Cilia were always defined by two markers-either two axoneme markers (Arl13b and polyglutamylated tubulin) or an axoneme and a basal body marker (Arl13b and gamma tubulin). SMB55 cilia are relatively short and so to avoid counting ciliary vesicles as cilia, only elongated Arl13b^+^ structures were counted. To quantify intensity of Smo at cilia, the cilium was defined by drawing a region of interest around the Arl13b^+^ region. Then average Smo pixel intensity in this region was measured. Smo intensity in a similarly-sized region of interest next to but not overlapping with each cilium was also measured and subtracted to normalize for background fluorescence. Similarly, Cep164 intensity was measured by drawing a region of interest around the gamma tubulin focus, measuring the average Cep164 intensity in this region, and subtracting background intensity of a nearby region of similar size. Cep164 was measured only at the Cep164^+^ centrosome (containing the mother centriole). If the mother centriole was not obvious (as in the case of Cep164 knockdown), the gamma tubulin focus with the higher Cep164 intensity was selected.

### Quantification and Statistical Analysis

#### Statistical analysis

All statistical analysis was performed in Prism 8 (Graphpad). Details for statistical tests used can be found in the figure legends. Figure legends also indicate the number of independent replicates performed and the number of cells analyzed for each condition of each replicate (n). All graphs show the mean and SEM. In all cases, significance was defined as \**p*-value ≤ 0.05, \*\**p*-value ≤ 0.01, \*\*\**p*-value ≤ 0.001, and \*\*\*\**p*-value ≤ 0.0001. ns indicates no significance.

#### Chi square analysis of sister cell fate

To test the null hypothesis that sister cells behave independently, we performed a Chi square analysis. All sister cell pairs from the live imaging data set in Figure 4C for which the cell cycle entry outcome was known for both sisters were included. For each time point (4-5 hr post SANT-1 addition, 6-7 hrs post SANT-1 addition, etc), the number of pairs observed to have each of 3 fates was counted: 1) both cells divided, 2) both cells arrested, or 3) the cells made different choices. Then, the “expected” distribution for the same number of pairs was determined by calculating the likelihood of each of the 3 fates if the two cells behaved independently based on the cell cycle entry frequency observed at that time point. These observed and expected distributions were then summed for all time points and compared using a Chi square analysis.

## End Matter

### Author Contributions and Notes

Conceptualization, E.K.H. and T.S.; Methodology, E.K.H.; Validation, E.K.H. and A.E.T.; Formal Analysis: E.K.H.; Investigation, E.K.H. and A.E.T.; Writing – Original Draft, E.K.H. and T.S; Writing – Review & Editing, E.K.H, A.E.T, and T.S.; Visualization, E.K.H.; Funding Acquisition, E.K.H., A.E.T., and T.S.; Supervision, T.S.

The authors declare no conflict of interest.

## Acknowledgments

We thank Rosalind Segal for the gift of SMB55 cells, James Ferrell for the gift of pTRIP eYFP-PCNA, Jamie Purzner and Alex Brown for advice regarding SMB55 cell culture, and all members of the Stearns lab for helpful discussion. This work was supported by a Ruth L. Kirschstein National Research Service Award Predoctoral Fellowship 1F31GM129950 to E.K.H., a Stanford Bio-X Undergraduate Research Fellowship to A.E.T., and National Institutes of Health grant 1R35GM130286 to T.S.

## Supplemental Information

**Figure S1:**
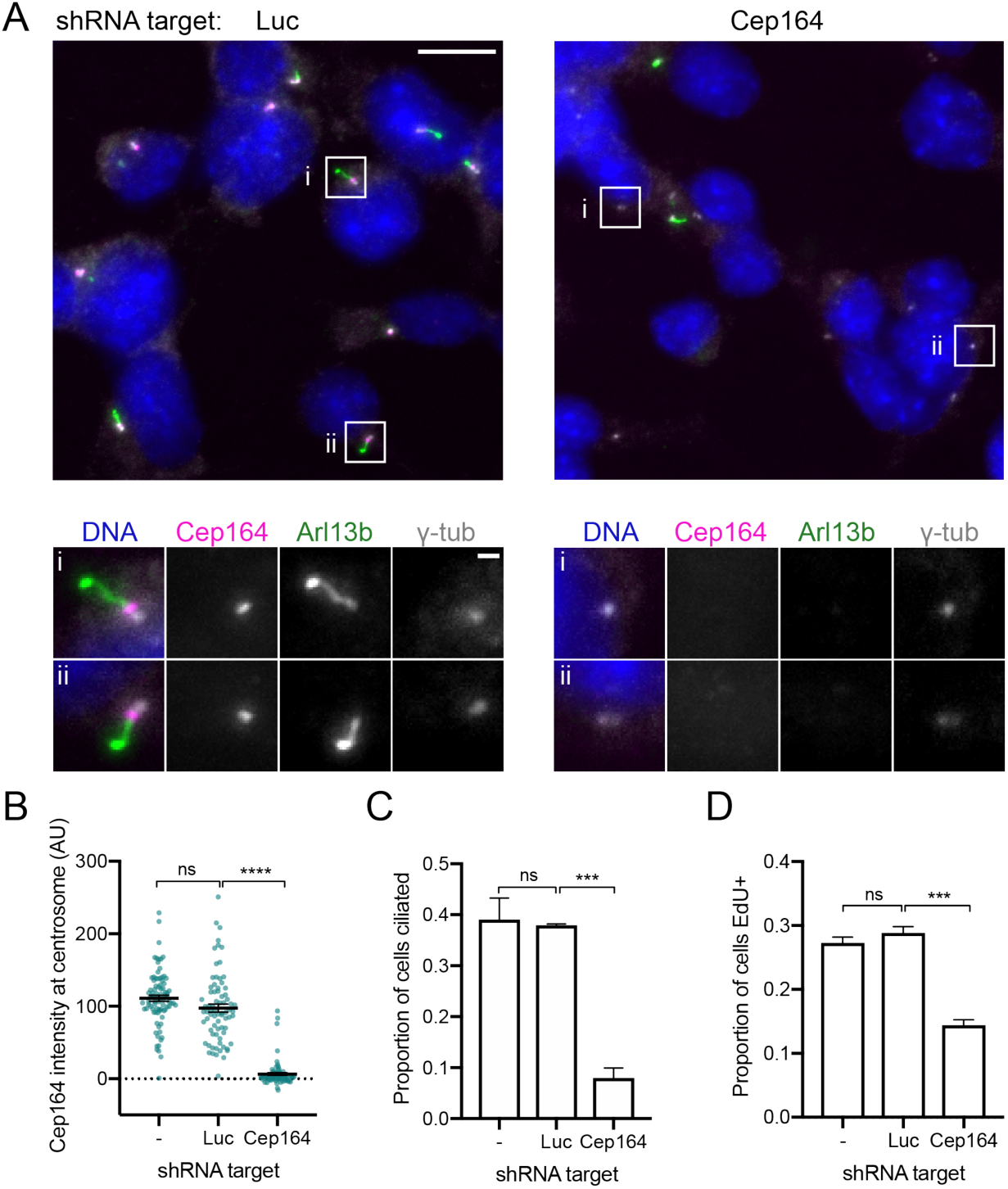
Primary cilia are required for SMB55 cell proliferation. Related to Figure 1. (A) Knockdown of Cep164 in SMB55 cells to confirm the requirement of primary cilia for SMB55 proliferation. SMB55 cells were lentivirally infected with shRNAs targeting Cep164, Firefly luciferase (Luc), or an empty vector (-) and assayed 4 days after infection by microscopy. We chose to knockdown Cep164, a distal appendage protein of the mother centriole required for membrane docking and ciliogenesis, because loss of Cep164 completely prevents cilia formation unlike other targets that shorten but do not fully eliminate the axoneme (Caspary et al., 2007; Graser et al., 2007; Schmidt et al., 2012; Wilson et al., 2012). Images show a reduction of Cep164 intensity at the centrosome (γ-tubulin) and cilia number (Arl13b) after 4 days of Cep164 knockdown. Scale bar: 10 μm, Inset:1 μm. (B) Cep164 intensity at the centrosome after Cep164 knockdown. Each point represents the measurement from a single cell. Mean±SEM from n≥75 centrosomes. Significance determined by one-way ANOVA and Tukey’s post hoc test. (C) Cilia number after Cep164 knockdown. Mean+SEM from 2 independent experiments, n≥200 cells in each. Significance determined by one-way ANOVA and Tukey’s post hoc test. (D) Proportion of SMB55 cells incorporating EdU during a 2-hour labeling 4 days after Cep164 knockdown. Cep164 knockdown significantly reduced EdU incorporation, showing the requirement of cilia for SMB55 cell proliferation. Mean+SEM from 3 independent experiments, n≥250 cells in each. Significance determined by one-way ANOVA and Tukey’s post hoc test.

**Figure S2:**
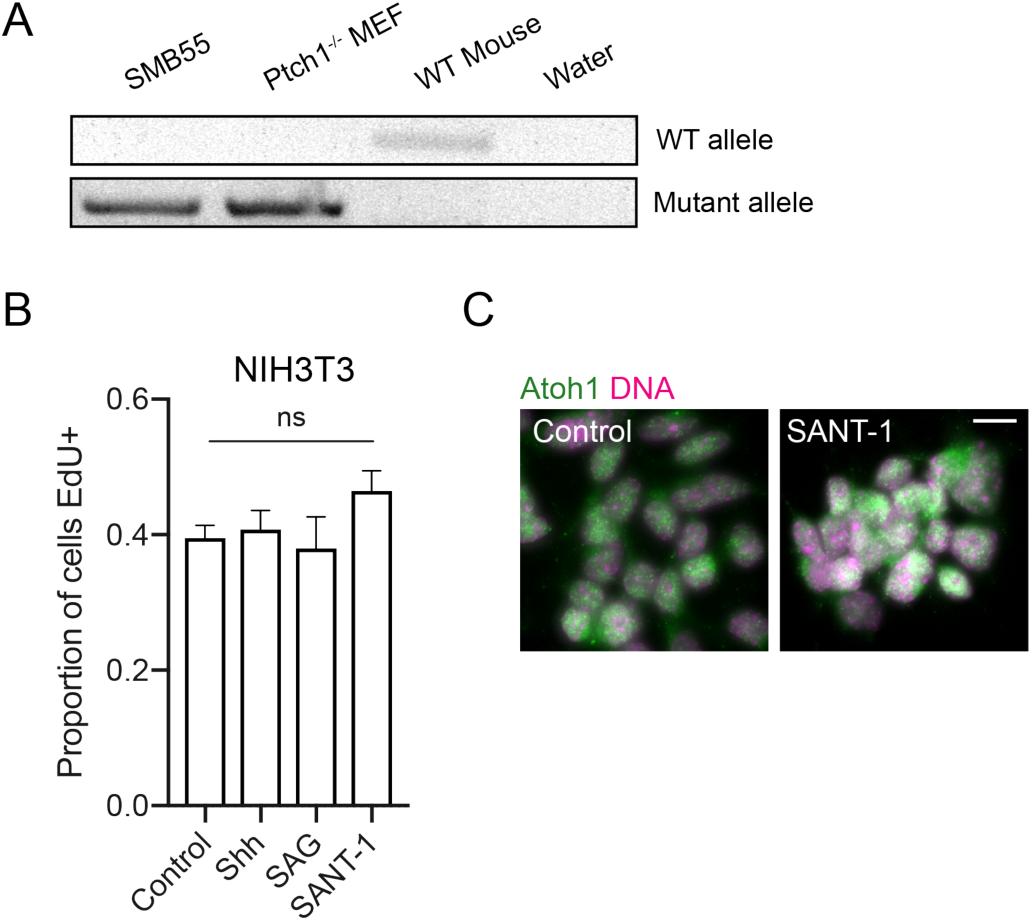
Hh pathway activity is required for *Ptch*^*-/-*^ SMB55 cell proliferation but not NIH3T3 cell proliferation. Related to Figure 1. A) Genotyping of SMB55 cells to test for presence of the mutant and wild-type *Ptch1* allele as described in (Goodrich et al., 1997). The wild-type allele was not present in SMB55 cells, confirming loss of heterozygosity at the *Ptch1* locus. *Ptch1*^*-/-*^ MEF and WT mouse genomic DNA were used as controls for the mutant and wild-type alleles respectively. (B) EdU incorporation in NIH3T3 cells in response to Hh pathway manipulation. NIH3T3 is a mouse fibroblast cell line that is highly responsive to Hh signaling but does not require Hh pathway activity for its proliferation. Graph shows the proportion of NIH3T3 cells incorporating EdU following 48-hour Hh pathway stimulation by 100 nM Shh or 100 nM SAG (Smo agonist), or inhibition by 100 nM SANT-1, and then a 2-hour EdU labeling. Mean+SEM from 3 independent experiments, n≥100 cells in each. Line over bars indicates no significant difference by one-way ANOVA. (C) Atoh1 expression in SMB55 cells after 48-hour SANT-1 treatment. Staining for Atoh1, a transcription factor and marker of cerebellar precursors, was used to assess differentiation state after SANT-1 treatment. The image shows nuclei from SANT-1 treated cells were Atoh1^+^, showing that Hh pathway inhibition does not cause differentiation within 48 hours. Scale bar: 10 μm

**Figure S3:**
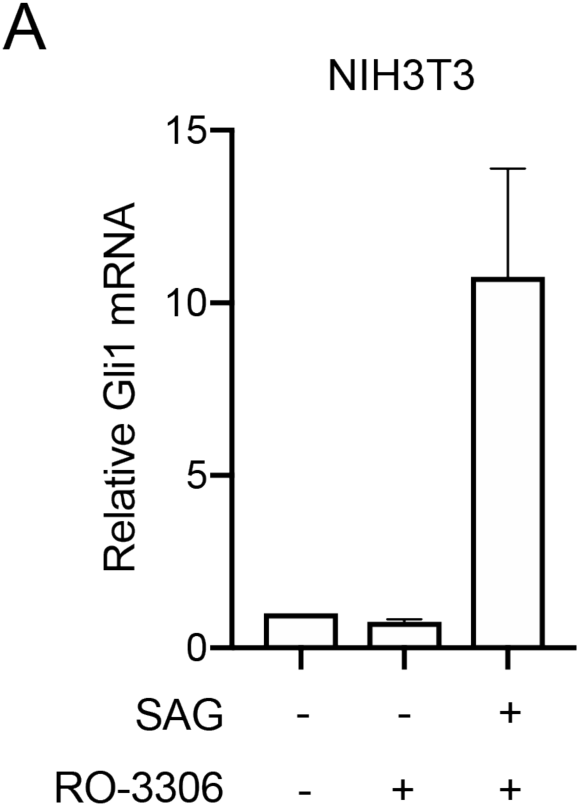
RO-3306 does not directly inhibit the Hh pathway. Related to Figure 3. (A) *Gli1* mRNA levels in NIH3T3 cells treated with RO-3306 and SAG. To decouple RO-3306’s effect on the cell cycle with any potential effect on the Hh pathway, NIH3T3 cells were grown to confluence and starved for 24 hours to induce cell cycle exit and ciliation prior to drug addition. Then these arrested cells were treated with SAG and RO-3306 for an additional 24 hours to induce pathway activity. Using RT-qPCR to measure *Gli1* mRNA, we found that RO-3306 treatment does not directly inhibit the Hh pathway. *Gli1* mRNA levels were normalized to GAPDH and expressed relative to the untreated controls. Mean+SEM, n=2

### Supplemental Video Legend

**Video S1. Example time-lapse imaging of SANT-1-treated SMB55 cells. Related to Figure 4.**

The timestamp (hh:mm) shows time since SANT-1 addition at 00:00. The white circles track two sister cells from an early M_1_ division that go on to divide again (M_2_). The daughter cells from these M_2_ divisions are marked by red circles and ultimately go out of frame. The yellow circles track two sister cells from a later M_1_ division (which have experienced longer pathway inhibition in the previous cell cycle) that do not divide again, persisting to the end of the image sequence.

